# Effective buprenorphine use and tapering strategies: Endorsements and insights by people in recovery from opioid use disorder on a Reddit forum

**DOI:** 10.1101/871608

**Authors:** Rachel L Graves, Abeed Sarker, Mohammed Ali Al-Garadi, Yuan-chi Yang, Jennifer S Love, Karen O’Connor, Graciela Gonzalez-Hernandez, Jeanmarie Perrone

**Affiliations:** Department of Emergency Medicine, University of Pennsylvania Health System, Philadelphia, PA; Department of Biomedical Informatics, School of Medicine, Emory University, Atlanta, GA; Department of Emergency Medicine, School of Medicine, Oregon Health and Science University, Portland, OR; Department of Biostatistics, Epidemiology and Informatics, Perelman School of Medicine, University of Pennsylvania, Philadelphia, PA

**Keywords:** Social media, opioid use disorder, reddit, buprenorphine, medication assisted treatment

## Abstract

Opioid use disorder (OUD) is a public health emergency in the United States. Over 47,000 overdose-related deaths in 2017 involved opioids. Medication-assisted treatment (MAT), in particular, buprenorphine and buprenorphine combination products such as Suboxone^®^, is the most effective, evidence-based treatment for OUD. However, there are a limited number of conclusive scientific studies that provide guidance to medical professionals about strategies for using buprenorphine to achieve stable recovery. In this study, we used data-driven natural language processing methods to mine a total of 16,146 posts about buprenorphine from 1933 unique users on the anonymous social network Reddit. Analysis of a sample of these posts showed that 74% of the posts described users’ personal experiences and that the top three topics included advice on using Suboxone^®^ (55.0%), Suboxone^®^ dosage information (35.5%) and information about Suboxone^®^ tapering (32.0%). Based on two models, one that incorporated ‘upvoting’ by other members and one that did not, we found that Reddit users reported more successful recovery with longer tapering schedules, particularly from 2.0 mg to 0.0 mg (median: 93 days; mean: 95 days), as compared to shorter tapering schedules investigated in past clinical trials. Diarrhea, insomnia, restlessness, and fatigue were commonly reported adverse events. Physical exercise, clonidine, and Imodium^®^ were frequently reported to help during the recovery process. Due to the difficulties of conducting longer-term clinical trials involving patients with OUD, clinicians should consider other information sources including peer discussions from the abundant, real-time information available on Reddit.

**Significance Statement:** Opioid use disorder (OUD) is a national crisis in the United States and buprenorphine is one of the most effective evidence-based treatments. However, few studies have explored successful strategies for using and tapering buprenorphine to achieve stable recovery, particularly due to the difficulties of conducting long-term studies involving patients with OUD. In this study, we show that discussions on the anonymous social network Reddit may be leveraged, via automatic text mining methods, to discover successful buprenorphine use and tapering strategies. We discovered that longer tapering schedules, compared to those investigated in past clinical trials, may lead to (self-reported) sustained recovery. Furthermore, Reddit posts also provide key information regarding buprenorphine withdrawal, cravings, adjunct medications for withdrawal symptoms and relapse prevention strategies.

## Background

Buprenorphine is one of the most effective evidence-based treatments for opioid use disorder (OUD). It is primarily prescribed as an oral or sublingual medication in office based primary care or addiction medicine practices, and it is increasingly being prescribed in hospitals and emergency departments to initiate therapy and bridge patients to outpatient programs. Although early treatment ambivalence and disengagement are not uncommon, once stabilized on buprenorphine, effective long-term recovery can be achieved. Despite the proven long term success in recovery with buprenorphine treatment, and advice from physicians to treat OUD as a chronic disease, many patients desire to taper or stop buprenorphine weeks, months or years into their recovery (1). Reasons for tapering and discontinuation may include stigma, expense, time, or the perception that non-abstinence-based addiction recovery is not valid (2, 3). Therefore, to meet patients’ preferences or treatment goals, it is important for prescribers to be knowledgeable about effective tapering strategies that are more likely to lead to successful discontinuation.

Although many patients attempt tapering, few studies have examined optimal tapering strategies. One study looked at 28-day vs 56-day tapering schedule in adolescents with OUD and found more opioid-free urines and more sustained retention in treatment with the longer tapering regimen (4). Another 12-year retrospective study of 1308 patients on buprenorphine showed that an estimated 15% tapered off buprenorphine, fewer than half of whom were medically supervised (5). Package insert information for the commonly used sublingual buprenorphine formulations do not include tapering advice and, in fact, may specifically advise against cutting, chewing, or swallowing the strips, all methods that patients structuring their own buprenorphine tapers commonly use (6). Physicians also prescribe film fractions for treatment and tapering (7), although there are currently no published best practice or guidelines regarding this practice.

Few studies have investigated providers’ or patients’ reasons for tapering or discontinuing buprenorphine. Studies have suggested that short term taper is not recommended as a stand-alone treatment (8) despite this common approach as medically supervised detoxification in many inpatient treatment facilities. One systematic review on this topic suggested that patients who discontinued buprenorphine therapy often did so involuntarily, and patients on lower buprenorphine maintenance at the time of tapering had better outcomes (2). Because of the unique pharmacology of buprenorphine and its formulation in sublingual filmstrips (*i.e*., Suboxone^®^), treatment centers, recovery groups, and social media channels have shared unique tapering strategies based on self-reported personal experiences. The information available through these less traditional sources has not been incorporated into conventional research on this topic. This is a missed opportunity given the large number of patients who structure their own tapers and/or taper without medical supervision. These sources have the potential to provide novel information regarding successful strategies for tapering buprenorphine, which may not be available from any other source. Due to the lack of evidence from clinical studies regarding effective tapering strategies, information curated through non-traditional sources from people who have tapered successfully may be useful for clinicians who prescribe buprenorphine. To obtain information directly from the consumers of buprenorphine, we focused on systematically analyzing relevant data from the social media forum Reddit. Reddit is a popular and rapidly-growing social networking website, with anonymity at the core of its design. As of July 2019, Reddit ranks as the fifth most visited website in the U.S. and 13th in the world, according to Alexa Internet and Wikipedia (9, 10). Recent advances in natural language processing (NLP), machine learning, and big data mining methods, coupled with the rise in mass social media adoption, have yielded novel information derived from social media in medical and public health research (11–13).

Using NLP, we mined and analyzed buprenorphine-related posts from the *subreddit* forum *|r|suboxone*, which allows its users to post anonymously. This anonymous network is rich in information about relatively sensitive and stigmatized topics, such as self-reported drug usage and recovery. Employing a data-driven, semi-automatic approach, we identified and quantified the topics discussed in this subreddit, which hosts all buprenorphine-related discussions, using the grounded theory approach (14). Within the forums, we found a high concentration of users who have personal experience (74.0% of posts) with opioid use disorder and provide support and advice to others on how to successfully use buprenorphine (55.0%), adjust buprenorphine dosage (35.5%) and taper buprenorphine (32.0%). We searched 16,146 posts from 1933 users for information about tapering strategies using NLP methods. We discovered elaborate descriptions of buprenorphine use and tapering strategies, including step-by-step guidelines from users who reported to have successfully recovered from OUD using buprenorphine. We manually analyzed and compiled these tapering strategies and summarized them in this paper. Notably, we found that longer tapering schedules, compared to those investigated in past clinical trials, were reported to be effective for sustained recovery, and that users found tapering down from 2.0 mg to 0.0 mg to be particularly challenging. We also characterized, qualitatively and quantitatively, strategies and substances that users described as helpful with recovery, and side effects encountered during and after terminating buprenorphine. In our quantification process, we utilized the ‘upvoting’ meta-information available from Reddit to scale information based on how many users had found the information useful.

## Results

### Data and Preliminary Analyses

In total, we mined 991 threads from the /r/suboxone subreddit containing 16,146 posts from 1933 users, including the original posts and comments (mean ~16.3 posts per thread; minimum number of posts = 1; maximum number of posts = 144). The earliest post was from February 2013 and the last post was from December 2018. The number of posts increased steadily over the years, with 32 posts in 2013, 142 in 2014, 595 in 2015, 1566 in 2016, 2352 in 2017 and 11,456 in 2018.

Our preliminary manual thematic analysis included 200 strategically selected posts from 10 threads (*Materials and Methods*) and verified the content-rich nature of the subreddit. This initial analysis revealed that 74.0% of the posts described personal experiences, 24.0% provided buprenorphine-specific information that were not personal experience, and 2.0% could not be categorized as either (*e.g*., speculations about intents of pharmaceutical companies). More fine-grained categorizations based on the grounded theory approach showed that the five most frequent topics of discussion included: advice on using buprenorphine for MAT (55.0%), information and guidance on buprenorphine dosage (35.5%), information about buprenorphine tapering (32.0%), side effects of buprenorphine and withdrawal information (30.0% combined), and specific questions about buprenorphine usage (21.0%). Table 1 presents all the subcategories and the distribution of these subcategories in the analyzed posts.

**Table 1.** Subcategories of topics of Reddit posts (n = 200) on the subreddit /r/suboxone and their distribution in the manually annotated data.

| SUBCATEGORY | CODE | DESCRIPTION | COUNT (% OF POSTS) |
| --- | --- | --- | --- |
| Cravings | c | discussions of cravings for other drugs or mode of administration (*e.g.* the need to inject) | 2 (1.0%) |
| Dosage | d | discussions of the buprenorphine dose users are taking, including taking too much or too little | 71 (35.5%) |
| Illegally obtaining, misusing, or abusing buprenorphine | ab | discussions about obtaining buprenorphine without a prescription or misusing buprenorphine | 5 (2.5%) |
| **Providing information and answering questions** | i | user-provided information about drugs/medications or answers to questions posted by other users | 42 (21.0%) |
| **Questions** | q | information seeking (e.g. how to come off buprenorphine) | 42 (21.0%) |
| **Relapse** | r | discussions or descriptions of relapse | 3 (1.5%) |
| **Side effects** | s | discussions of side effects of buprenorphine use | 27 (13.5%) |
| **Withdrawal** | w | discussions or descriptions of withdrawal from buprenorphine | 33 (16.5%) |
| **Using other drugs to help with withdrawal** | u | use of another drug for withdrawal (with the drug used specified) | 3 (1.5%) |
| **Use of other substance(s) to mediate symptoms of opioid withdrawal** | os | discussion or description of substance(s) other than buprenorphine (e.g., kratom, loperamide etc.) for mediating symptoms of opioid withdrawal | 15 (7.5%) |
| **Specific advice on how to use buprenorphine** | as | user-provided advice on how to use buprenorphine | 110 (55.0%) |
| **Advice on something other than how to use buprenorphine** | ao | user-provided advice not related to buprenorphine use | 28 (14.0%) |
| **Stigma** | st | expression or description of stigma associated with buprenorphine use | 3 (1.5%) |
| **Tapering** | t | discussion of strategies or experience with lowering buprenorphine dose over time | 64 (32.0%) |
| **Other** | o | content outside of categories above | 4 (2.0%) |

### Tapering Strategies

While many topics were discussed in the /r/suboxone subreddit forum, the most unique information we obtained were self-reports of detailed tapering strategies for buprenorphine. People who had successfully used buprenorphine to treat their OUD provided detailed information and tapering recommendations to others. Patients who had been taking Suboxone^®^ described specific tapering strategies that involved cutting 2 mg strips into as many as 32 pieces and taking decreasing doses over time before terminating or ‘jumping off’.^1^ The same principle applies to users taking other buprenorphine-containing products. Reddit users who had been taking Subutex^®2^ described methods for dissolving solid pills in water and decreasing their intake volume over time in order to taper. Based on the various descriptions of methods used for tapering in our preliminary dataset, we developed natural language expressions to identify posts from the entire dataset which could potentially provide complete tapering schedules, including dosages and durations. We discovered that compared to the typical smallest doses of buprenorphine recommended in the published literature, self-reports of successful tapering involved much smaller doses. The most frequent/popular successful final dose before termination was 0.063 mg; 0.063 mg and 0.125 mg were both popular doses because 2 mg strips of Suboxone^®^ can be cut into 16 equal pieces so that each piece contains approximately 0.125 mg, or into 32 pieces so that each piece contains approximately 0.063 mg.

The design of Reddit enables users to upvote or downvote (analogous to Facebook likes) posts based on their utility. To quantify successful tapering strategies and terminal doses of buprenorphine from Reddit, we used two models: one that included the number of posts that presented information about (i) successful tapers and (ii) a specific dose at which the user quit (*unscaled model*); and a model that scaled the buprenorphine dose frequencies found in the first model by the number of upvotes they received (*scaled model*). Figure 1 presents the frequencies of the buprenorphine doses from both the unscaled (top) and scaled (middle) models, and the relative frequencies obtained by the two models for comparison. In both models, 0.063 mg was the most popular quitting dose, followed by 0.125 mg. Comparison of the relative dose frequencies of the two models shows that when upvoting information is incorporated, the popularities of smaller doses are further amplified, as both the 0.063 mg and 0.125 mg doses have higher relative frequencies in the scaled model than the unscaled model. The highest successful quitting dose reported was 2 mg, which only received 2 upvotes, indicating that other users may not have found this information to be useful.

**Figure 1.**
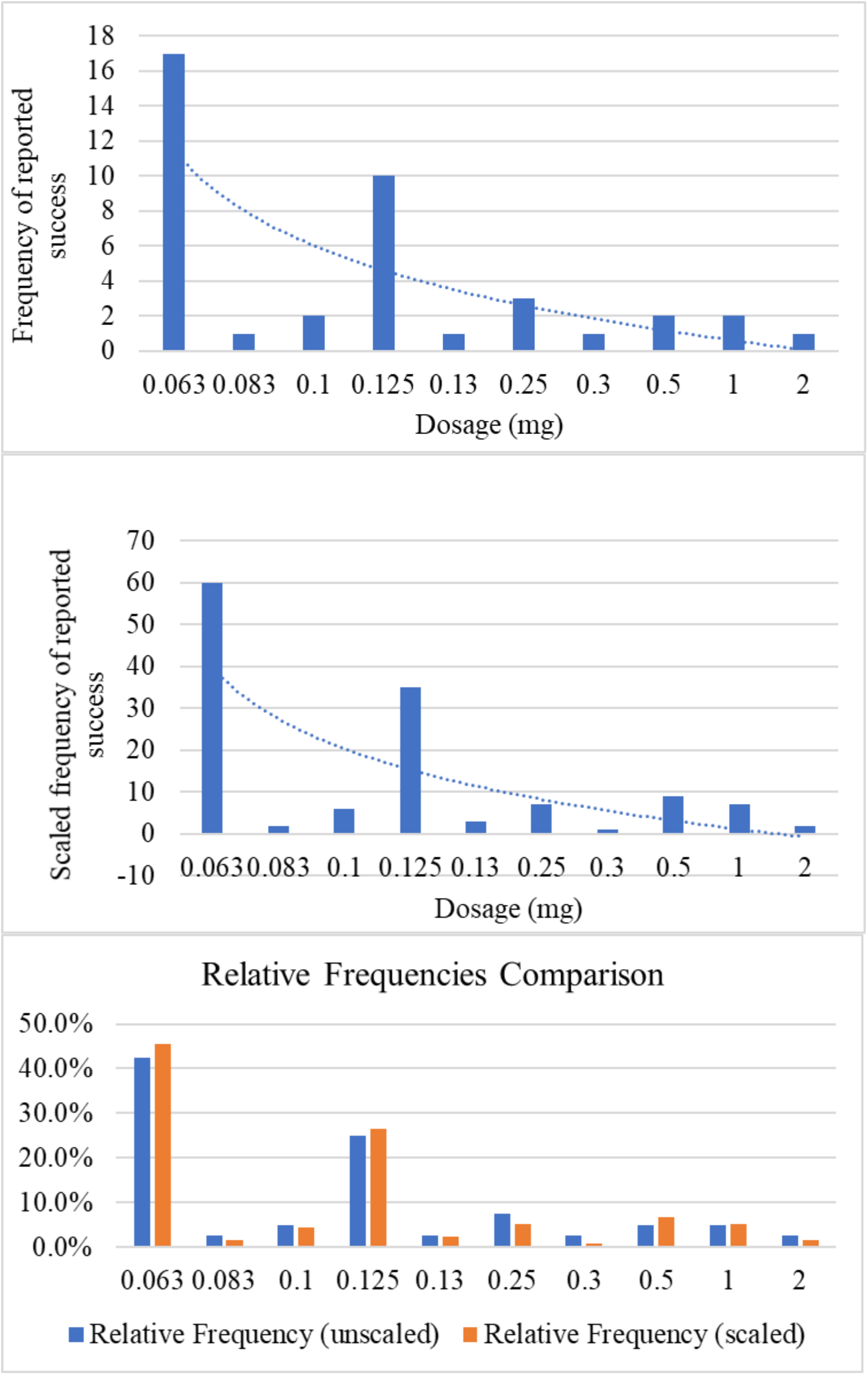
Frequencies of dosages (mg) at which the selected subset of Reddit users reported to have ‘jumped off’ Suboxone.

23 users provided sufficient information about their tapering schedules, including dosages and durations at each dose, for us to quantify (in some cases approximately) the number of days they took to taper from 2 mg to 0 mg. Figure 2 illustrates the tapering durations via a boxplot. Most described schedules were between 60-120 days, with a median of 93 days. The figure also shows 3 outliers, which are users who tapered over longer durations compared to the others—beyond 1.5 times the inter-quartile range from the upper quartile. For 19 users, we could determine or estimate their specific tapering schedule based on their posts. Figure 3 illustrates these tapering schedules using a multi-line graph, showing up to 180 days before the final dose/last day. In the figure, the thickness of the lines is proportionate to the number of users reporting the specific schedule.

**Figure 2.**
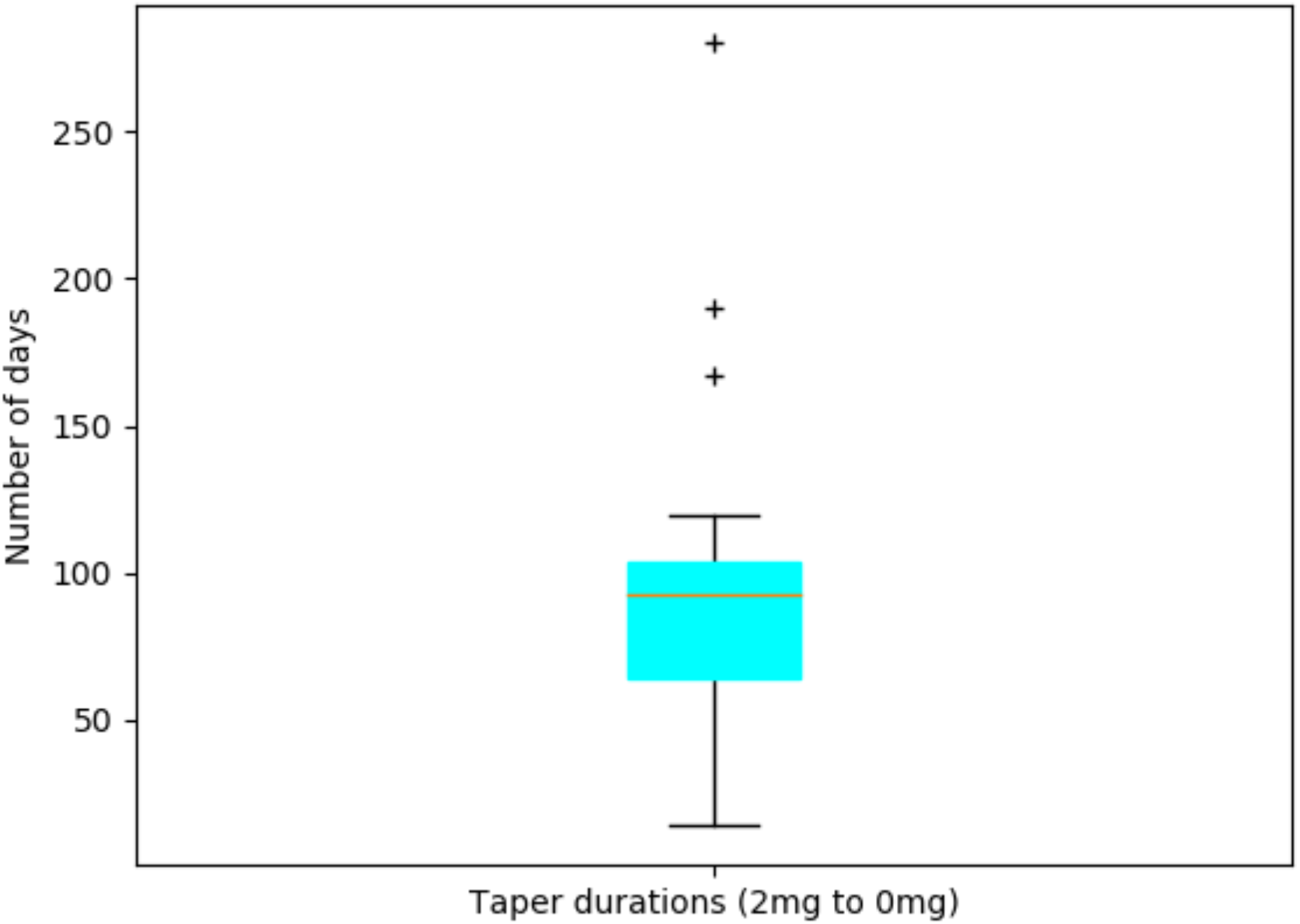
Tapering durations in days from 2mg to 0mg for the subset of Reddit users for whom it was possible to curate this information (n=23).

**Figure 3.**
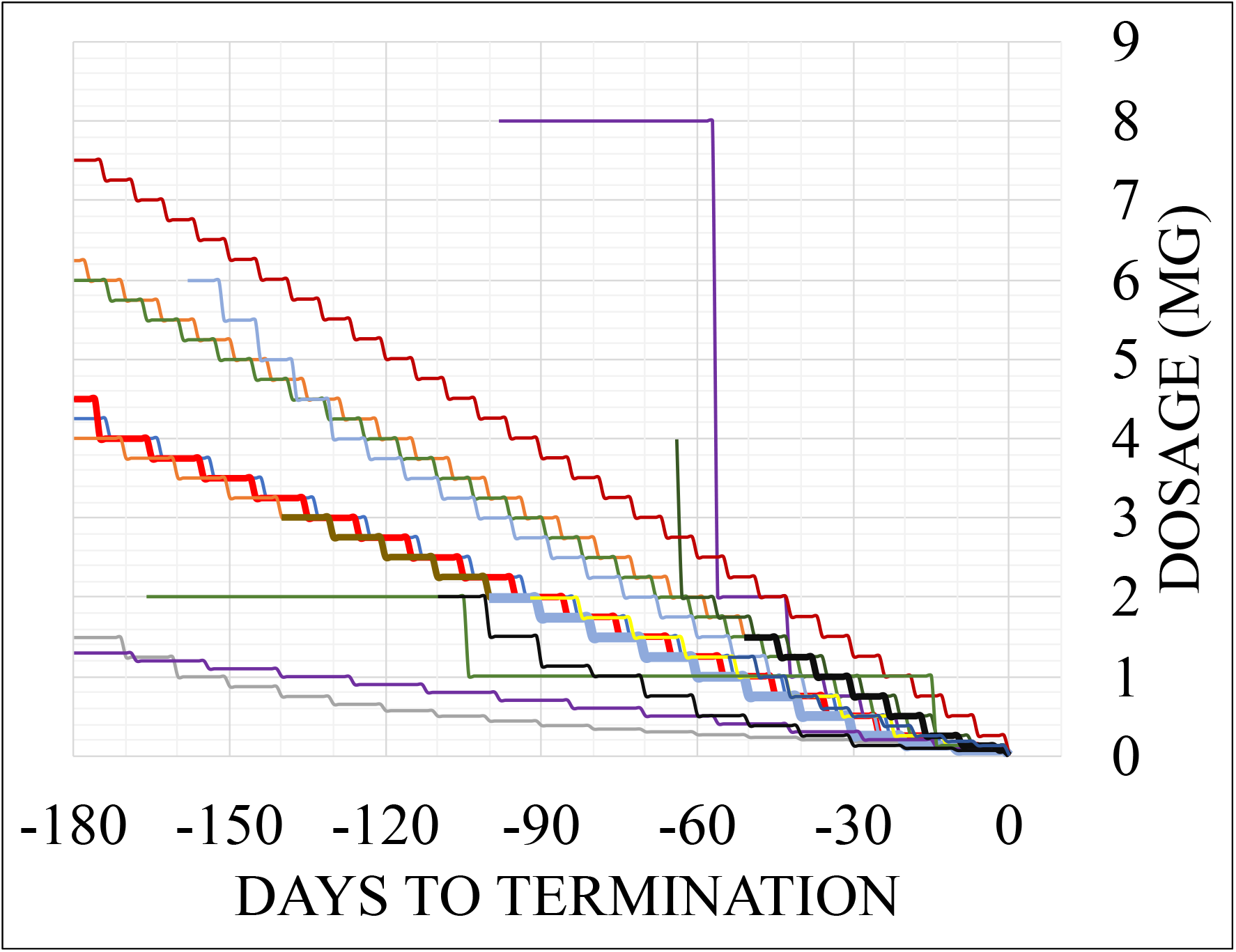
Tapering schedules for up to the last 180 days. Thicker lines indicate more users reporting the same schedule.

### Additional Buprenorphine MAT-related Information

For Reddit users who describe successful use of buprenorphine therapy to treat OUD and present information about their tapering strategies, we attempted to quantify (i) any other substances or methods they reported to be useful and (ii) the side effects or adverse events they reported. Like the previously discussed analyses of tapering doses, we used two models for each of these, one unscaled and the other scaled by the number of upvotes.

Figure 4a shows the relative frequencies of various adverse events in the unscaled and scaled models. Diarrhea, insomnia, fatigue and restless leg syndrome (RLS) appear to be the most common adverse events, with the relative frequency of insomnia being much higher in the scaled model as compared to the unscaled model. Conversely, the relative frequencies of fatigue and diarrhea were lower in the scaled model.

**Figure 4.**
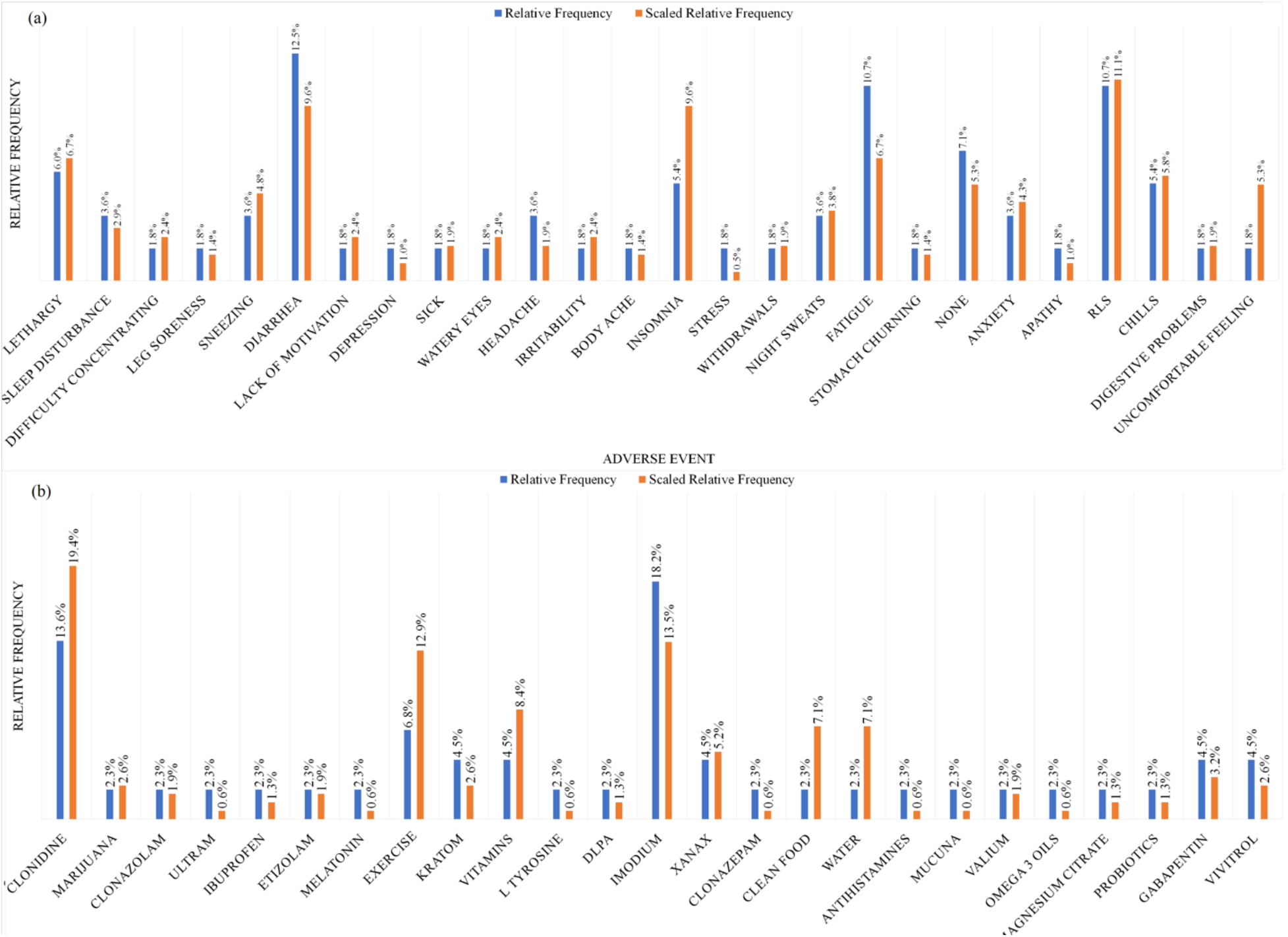
(a) Relative frequencies of adverse events reported after tapering suboxone by the Reddit cohort studied (n=50). ‘none’ represents users who specifically reported not needing or using any other substance or method. (b) Relative frequencies of substances and other methods (e.g., exercise) reported by the Reddit cohort to be helpful in dealing with withdrawals and other symptoms experienced after quitting suboxone (n=50).

Figure 4b presents the relative frequencies of the substances and/or methods used by users who successfully treated OUD in the unscaled and scaled models. Substances or methods that received more upvotes had higher relative frequencies in the scaled model compared to the unscaled model. Clonidine and Imodium^®^ (loperamide) are commonly mentioned, with clonidine having a much higher relative frequency in the scaled model and loperamide a much lower relative frequency in the scaled model as compared to the unscaled model. Exercise was reported to work well, with particularly high relative frequency in the scaled model, meaning that it received many upvotes.

## Discussion

Our study shows that Reddit specifically, and social media in general, may provide important information to clinicians and caregivers who treat patients suffering from OUD. OUD sufferers represent a difficult population for conducting clinical trials and studies as they are difficult to follow, perhaps particularly if they relapse. Therefore, the information available from Reddit may not actually be available via other sources. From the discussions we found on the /r/suboxone subreddit, it was evident that many people were seeking support and advice from their peers, and the detailed nature of many posts suggest a level of comfort that users have in anonymously sharing their personal experiences. Although it is not clinically recommended, many OUD sufferers choose to taper buprenorphine over time, with the view of terminating its use at some point. Therefore, to adhere to the principles of patient-oriented care, clinicians need to provide treatment plans that maximize the chances of patients’ successful recovery.

In addition to the information-rich nature of the discussions, a few discoveries we made in this study stand out, which are summarized in the following list:

1. Long tapering durations, compared to those tested in past clinical studies, are more likely to lead to successful recovery.
2. As suggested by the outliers in Figure 2, some users may need to taper over much longer durations than others to successfully recover.
3. The most difficult part of tapering is going from 2mg to 0mg.
4. Common side effects encountered by patients during their effort to taper include insomnia, RLS, diarrhea, fatigue and lethargy (Figure 4a).
5. Certain medications and physical exercise (Figure 4b) may help patients cope with some of the side effects encountered during tapering.

### Relevance of Social Media

Social media adoption has accelerated rapidly over the last decade, and the use of social media data for health-related research has steadily increased in recent years. Data from the PEW Research Center shows that as of February 2019, 69.0% of U.S. adults use Facebook, 22.0% use Twitter, and 11.0% use Reddit, and 22.0% of younger users (ages 18 to 29) use Reddit (15). Social media provides researchers access to knowledge posted directly by users in near real time. Social media data have been utilized for infectious disease outbreak monitoring (16–18), adverse drug reaction detection (19, 20), understanding behavioral health patterns (*e.g*., predicting depression) (21, 22), and for characterizing prescription medication abuse/misuse (23–25).

Social media has also been used for studying buprenorphine self-management. A study published in 2017 (26) showed that online discussion board posts contain information about Suboxone use and self-treatment. The study suggested that the users who posted in the discussion boards seemed to trust their peers more than pharmacists/physicians. Self-management activities cited in the paper included altering the amount of Suboxone taken to modify or changing the administration route. The authors concluded that information about shared knowledge and behaviors of people with substance use disorders are important to healthcare providers because of the previously unknown precautions and risks associated with self-treatment. Another study analyzed 100 Reddit posts and categorized them according to the number of criteria for opioid use disorder as outlined by the Diagnostic and Statistical Manual of Mental Disorders (DSM-5) described in each (27). This study showed that 33% of the users described enough symptoms in their posts to meet DSM-5 criteria for OUD. Many users sought advice (43.0%) or support (24.0%) from their peers, and 95% of the posts contained at least 1 therapeutic factor.

Other studies relevant to our work include those that describe non-medical prescription medication use, and those that describe opioid misuse or abuse. Chan *et al*. (28) manually characterized Twitter posts mentioning opioids and showed that among tweets mentioning personal use of opioids, the vast majority (~88.0%) represented non-medical use. Graves *at al*. (29) and Chary *et al*. (30) used unsupervised machine learning methods (*i.e*., without manually annotated training data) to derive meanings from large, unlabeled sets of tweets mentioning opioids and found correlations between numbers of tweets and state-level information from the National Surveys on Drug Usage and Health (NSDUH) and county-level opioid death rates. In our past work on the topic, we showed that supervised machine learning is capable of automatically characterizing abuse-related discussion on Twitter and that the temporal patterns of abuse detected from social media convey information identical to that obtained from prior manual analyses (23). While earlier research efforts have established the utility of social media for studying drug abuse/misuse, addiction and recovery, there is no study that has attempted to mine and quantify strategies for self-driven MAT from social media. As methods for mining knowledge from social media are maturing, we are gradually beginning to see the true value and possible applications of knowledge garnered from social media. Most envisioned applications of social media mining and research advances to date, including our own past research, have been in the population/public health realm (31–34). However, perhaps due to the relatively low evolution and translation of social media research into public health or medical practice, some studies have raised questions about the real-life effectiveness of social media (35, 36). The study described in this paper illustrates an interesting, unique and useful application social media-based knowledge mining that can inform clinicians who are treating patients with opioid use disorder.

Among the various social media platforms that have been studied, Reddit has structural characteristics that make its content particularly compelling for studies similar to ours. Reddit uniquely combines the ability to interact as a completely anonymized user with access to large populations of people who are interested in or have experience in a particular topic. In essence, Reddit is a forum constructed around topics rather than around the user. It is not a forum for self-promotion, image-branding, or other individual-oriented metrics. Content is upvoted or downvoted on its perceived merit, separated by design from the identity of the user to a greater extent than other popular social media platforms. This “idea-centric” model differs from the other social media platforms the primary purpose of which is to project or promote an individual or institution. Our findings will inform buprenorphine prescribers to help guide patients in optimizing medication and more insights into discontinuation strategies to sustain recovery from OUD.

### Limitations

Reddit users are disproportionately likely to be younger, male, and Hispanic, and may not comprise a representative sample of the United States population. Comments about buprenorphine products, tapering schedules, withdrawal, or other topics may not be made by the individuals with personal experience, so content analysis may reflect public perspective or hearsay rather than reliable peer information. Reddit posts are anonymous, so it is difficult to follow timelines or outcomes of suggested strategies. Posts are also based on self-reporting of Reddit users’ perspective or experience rather than on objective data (*e.g*., documentation of drug-free urine samples in order to determine abstinence from opioid use).

### Summary

Content about Suboxone from the social network Reddit may be a resource for patients and providers to gain information and insight into Suboxone therapy. Specific themes around Suboxone tapering strategies may be valuable due to a gap in the medical literature on this topic. Use of Suboxone among patients as reflected on Reddit may be inconsistent with current medical practice and introduce areas for improved practice and further exploration.

## Methods

### Study Design

This was a mixed methods observational study of posts from the /r/suboxone subreddit of Reddit. The study protocol was approved by the University of Pennsylvania Institutional Review Board (IRB). Our initial exploration, which was performed by browsing the social network website, revealed that this chosen subreddit provided the platform for discussion related to buprenorphine.^3^ All the data used was publicly available at the time of the study, so informed consent from the users were not obtained. The study did not involve any interaction with the Reddit users.

### Data Collection Strategies for Initial Analyses

Reddit is one of the most popular and heavily-used social media platforms, and it enables users to post and discuss information anonymously in topic-specific forums (called *subreddits*). In July 2019, there were almost 1.7 billion visits to Reddit, and it currently hosts over 130,000 sub-forums and communities. Reddit ranks among the most popular social networks in the US with over 26 million logged-in users, with 15% of adult males and 8% of adult females in the US actively using the social network.^4^ Providing a safe and anonymous space for users is at the core of Reddit’s design and culture, with elaborate community-established rules and etiquettes known as ‘*rediquette*’.^5^ Due to the emphasis on anonymity of this platform, it is particularly popular for openly discussing sensitive and stigmatized topics, such as opioid addiction and recovery.

We collected data from Reddit using the PRAW API.^6^ All available posts from the /r/suboxone subreddit were collected. The research and analyses we present here were data-driven (as opposed to hypothesis-driven), reliant on the information shared by the community members. Due to the lack of past research and data sources on this topic, a data-driven research approach was essential—we needed to first identify and quantify the types of information available from this network, and then devise strategies for mining the relevant information. We employed NLP and social media text mining methods to collect and curate the buprenorphine-related conversations, followed by qualitative and quantitative analyses to summarize the pertinent findings. To better gauge the contents of the /r/suboxone subreddit, we first conducted a preliminary study involving 200 posts from 10 threads within the subreddit (37).

Since it is impossible to manually analyze all the posts within the /r/suboxone subreddit, we devised a strategy to include only *useful* or *popular* posts and to sample posts from a variety of users, rather than many posts from the same user. To determine usefulness/popularity, we used the voting feature of Reddit. Reddit allows users to upvote or downvote posts, and the aggregate of the votes (henceforth: score) can be used as an indicator of usefulness. For the analyses of 200 posts, we only included posts that have a net score of +3, indicating that at least 3 users found the post to be informative or useful. Since it is possible for the same user to be repeatedly posting in the subreddit, we included a maximum of 3 posts by any user.

### Annotation Categories and Manual Categorization

Using the grounded theory approach (14), posts were first categorized into fine-grained topics (*e.g*., withdrawal, stigma, relapse etc.). These were then merged into coarse grained topics. We prepared thorough annotation guidelines based on the contents of a sample of annotated tweets (see supplement), which were followed during the manual categorizations. Our final annotations contained three coarse-grained/high-level categories: *information* (i), *personal experience* (p), and *other* (o), and 15 fine grained topics (14 + other), as shown in Table 1. The fine-grained categorization was a multi-label classification, with each post labeled as at least one category or ‘other’.

Due to the structure of Reddit threads, individual posts often lack context. For example, a post that meets both of the above criteria may not be the original post that starts a thread but rather a response to another post. Therefore, during manual annotation, we did not annotate individual posts. Instead, for each post selected, we also included its parent post and annotated the pair. Table 2 presents a simplified and short example of how the data were presented for manual annotation/categorization.

**Table 2.** Simplified sample of how the data was presented during manual annotation, illustrating the pairing of comments with a parent comment and a child comment.

| Post Title | Parent Comment | Coarse Category | Fine Category | Upvotes* | Child Comment | Coarse Category | Fine Category |
| --- | --- | --- | --- | --- | --- | --- | --- |
| Jumped to 0 today | I have been on subs for over 5 years, not prescribed but a connect who is. I have quit once before but I stopped using all drugs last time for about 2 months (for a job). Man am I feeling horrible today. I jumped from about a 0.0625 mg which is pretty much nothing. I have been taking cbd and its been helping me sleep, I also have some kratom in the back pocket just in case. But I dont know if I want to use it. Any thoughts/ideas would be greatly appreciated. | p | as, ao, d, t | 4 | Congrats man I'm wishing you well. I don't have any advice as i'm just getting started with subs really. That is a super low dose I'm curious how long did it take to get to the 0.0625? How do you measure that, and did you even feel anything off that? Id imagine if you've been on that dose for awhile the withdrawal won't last too long or be that bad as getting down to that dose would have been the hard part. A lot of it has to be mental at that point but I know that can be the worst part and can be more challenging then the physical. Have you ever tried gabapentin? Its the only thing that ever worked for my wd's of dope even worked way better then kratom but might not be worthwhile here. Good luck man! | p | as, t, w, ao, d |

\*Upvotes counted at the time of collection

### Data Selection for Studying Tapering

We used the posts labeled as *t* in our initial sample to collect more posts about tapering for detailed analyses. We identified keywords and expressions that were particularly associated with detailed tapering-related information. These expressions included the words and phrases that indicated taper or quitting (‘*taper*’, ‘*jump off*’, ‘*jumped*’, ‘*quit*’); terms that indicated durations (‘*week(s)*’, ‘*month(s)*’, ‘*year(s)*’, ‘*day(s)*’); and terms indicating doses (‘*mg*’, ‘*mils*’, ‘*milligrams*’, and numbers with decimal places *e.g*., ‘*0.5*’). For a post to be included for detailed analysis, it had to match at least two of these three expressions. In addition, since posts on Reddit frequently contain misspellings, inexact references and colloquial expressions, we employed a *fuzzy* searching strategy—we passed the text through word windows of size 3 and computed their Levenshtein ratios with our pre-curated expressions. Posts that had at least a maximum Levenshtein ratio of 0.90 were included. Finally, we randomly sampled the retrieved posts for manual analysis.

### Data Selection for Studying Side Effects and Other Substances/Methods

Since we only wanted to study side effects for those users who to have successfully recovered, we only computed these frequencies after completion of our tapering analysis. These concepts were detected and grouped manually. Distinct lexical representations of the same concept (*e.g*., ‘*RLS*’ and ‘*restless legs*’) were grouped before computing frequencies. The relative frequencies of the terms are computed using the formula: 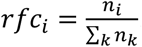, where *n_i_* is the number of occurrences of concept *i* and 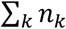 is the sum of occurrences of all concepts in the set.

## ACKNOWLEDGMENTS

Research reported in this publication was supported by the National Institute on Drug Abuse of the National Institutes of Health under Award Number R01DA046619. The content is solely the responsibility of the authors and does not necessarily represent the official views of the National Institutes of Health.

## Footnotes

1 *‘jumping off* is a phrase that is commonly used by the subreddit community to describe stopping using buprenorphine.

2 Subutex^®^ is the Canadian brand name for a buprenorphine formulation that does not contain naloxone in contrast to Suboxone^®^ which is the US brand name for a buprenorphine/naloxone combination product.

3 Suboxone^®^ is the tradename of the sublingual film containing buprenorphine and naloxone.

4 Statistics available from: Statista. https://www.statista.com/statistics/443332/reddit-monthly-visitors/ [Accessed 11/05/2019].

5 Full details of Reddiquette available at: https://www.reddit.com/wiki/reddiquette. Updated: 2018. [Accessed 11/06/2019].

6 Available at: https://praw.readthedocs.io/en/latest. [Accessed 07/16/2019].

## Notes

#### Summary of Updates

Acknowledgment added. Minor corrections made to the writing.

